# Critical Period Plasticity is Associated with Resilience to Short Unpredictable Stress

**DOI:** 10.1101/2025.02.26.640466

**Authors:** Robert Williams, Charlie Van Den Oord, Erica N. Lee, Samuel C. Fedde, Gia L. Oscherwitz, Adema Ribic

**Author notes:** Equal contribution (Robert Williams, Charlie Van Den Oord, Erica N. Lee).

## Abstract

Low resilience to stressful events can increase the risk of anxiety and depression. Resilience decreases with age, parallel to drastic changes in the quality of brain plasticity from juvenile to old age, suggesting that the type of plasticity found in the maturing brain promotes resilience. To indirectly test this, we administered short unpredictable stress to adult male and female wild type (WT) C57BL/6 mice, as well as to two groups of mice characterized by heightened cortical plasticity: adolescent C57BL/6 WT mice and adult mice that lack SynCAM 1 (Synaptic Cell Adhesion Molecule 1), a critical plasticity brake in the mature brain. We found that short unpredictable stress robustly increased core body temperature in all groups of mice, indicative of stress-induced hyperthermia (SIH) and confirming the efficacy of the stress paradigm. However, depressive-like behavior as measured though tail suspension test was increased in adult WT mice only, supporting that the type of plasticity found in the immature brains of adolescent WT and adult SynCAM 1 knockout (KO) mice promotes resilience to stress. All three groups of mice showed a mild increase in locomotor activity after stress, suggesting that the quality of plasticity does not correlate with resilience to anxiety-like phenotypes. Our study hence provides indirect evidence for the protective role of developmental plasticity during stress and points to new mechanisms that promote resilience to stress-induced depression.

## Introduction

Stressful life events are strongly linked to the onset of depression (McEwen, 2012; McEwen et al., 2015; Czéh et al., 2016), which is one of the leading causes of disability in the USA (WHO). Yet, stress does not increase the risk of depression in individuals that are resilient: capable of successfully adapting to stress exposure (Faye et al., 2018; McEwen and Morrison, 2013; Russo et al., 2012). Stress resilience is thought to decline with age, parallel to changes in the quality of brain plasticity, which transitions from robust critical period plasticity during in the developing brain to a restricted, top-down modulated process in the adult brain (Ribic, 2020). While the link between plasticity and resilience is still unclear, a previous study demonstrated that chronic restraint stress results in a reversible reduction in the length of dendritic branches in the prefrontal cortex (PFC) of young, but not aged rats, in whom the reduction persists (Bloss et al., 2010). Heightened brain plasticity during postnatal development, as well as interventions that change the quality of plasticity in adulthood, can hence promote resilience to stress (McEwen, 2016, 2012). In support of this notion, antidepressants markedly increase plasticity in the adult brain and change its quality to a more juvenile/adolescent-like (Maya Vetencourt et al., 2008; Phoumthipphavong et al., 2016). Environmental enrichment, a classical behavioral paradigm that also “rejuvenates” the adult cortex (Greifzu et al., 2016, 2014), mitigates the effects of stress across the lifespan (Macartney et al., 2022; Lehmann and Herkenham, 2011; Hegde et al., 2020). However, both antidepressants and environmental enrichment have systemic effects on the brain, and it is still unclear if they promote resilience and mitigate the effects of stress through altered quality of brain plasticity.

In mice, a species of choice for genetic manipulations of plasticity, stress during late postnatal development has adverse effects on adult behavior and physiology (Peleg□Raibstein and Feldon, 2011; Brydges et al., 2014). However, stress administered during the juvenile period does not negatively impact behavior during adolescence, a period in which plasticity is still heightened, supporting the protective role of developmental plasticity during periods of stress (Peleg□Raibstein and Feldon, 2011). Interestingly, genetic mouse models in which developmental windows of plasticity do not close show altered behavioral responses to adverse experience (Ribic, 2020), from impairments in fear learning (Park et al., 2016; Banerjee et al., 2017) and juvenile-like responses during fear conditioning paradigms (Bhagat et al., 2016; Gogolla et al., 2009), to facilitated fear erasure (Yang et al., 2016) and fear-conditioned response (Miwa et al., 2006). While these studies indicate a complex association between plasticity and adverse experiences, they also point to a protective role of critical period plasticity after fear-inducing events and suggest that reintroducing juvenile/adolescent-like plasticity in adult brain could promote stress resilience.

To begin to address this, we tested the depressive-like and anxiety-like behaviors after a week-long bout of unpredictable stress in mice during early adolescence (postnatal days 28-35), an age at which cortical areas still display critical period plasticity (Ribic, 2020). We further tested if adult mice with extended critical period plasticity through deletion of SynCAM 1 (Synaptic Cell Adhesion Molecule 1)(Ribic et al., 2019) respond to stress in a manner similar to adolescent mice. SynCAM 1 is an immunoglobulin domain-containing Type I transmembrane protein that organizes excitatory synapses (Perez de Arce et al. 2015). SynCAM 1 protein is expressed throughout the brain, with expression gradually increasing during postnatal development and plateauing in adulthood (Ribic et al. 2019; Robbins et al. 2010; Thomas et al. 2008; Fogel et al. 2007). Constitutive loss of SynCAM 1 enhances hippocampal long-term depression *ex vivo* (Robbins et al. 2010) and, much like antidepressants and environmental enrichment, shifts the quality of *in vivo* plasticity in the adult visual cortex to a juvenile/adolescent-like state (Ribic et al. 2019). Further, adult SynCAM 1 knockout (KO) are resilient to fear conditioning (Park et al. 2016), suggesting that loss of SynCAM 1 plays a protective role during traumatic experiences.

Using a battery of physiological and behavioral assays, in this study we describe the effects of short-term unpredictable stress on the physiology, as well as depressive-like and anxiety-like behaviors of young and adult wild type (WT) male and female C57BL/6 mice and adult male and female SynCAM 1 KO mice, to test the association between the quality of plasticity (adult vs juvenile/adolescent like) and behavioral resilience to stress.

## Methods

*Mice*. All mice were maintained on C57BL/6 background (The Jackson Laboratory, Bar Harbor, ME) on a reverse 12:12 light:dark cycle, with food and water *ad libitum*. Young animals from both sexes were used during the 4^th^ and 5^th^ week after birth. Adult animals from both sexes were used from 2-4 months of age. SynCAM 1 KO mice (Fujita et al., 2006) and their WT littermates were maintained on a C57BL/6 background and used at 2-4 months of age. Animals were randomly assigned to experimental groups and littermates were group housed for the duration of experiments, except during social isolation. Animals from multiple litters were included in the study to avoid any litter effects. Animals were treated in accordance with the University of Virginia Institutional Animal Care and Use Committee guidelines.

### Unpredictable stress

Mice underwent unpredictable stress over a course of one week, which consisted of the following stressors: physical restraint in a 50 ml falcon tube for adult mice or 20 ml plastic scintillation vial for young mice (3 times a week for 1 hr), placement in a cage with no bedding (2 times a week for 2 hrs) or wet bedding (250 ml of distilled water added to a standard cage with bedding 2 times a week for 2 hrs), tilting the cage at a 20º angle (2 times a week for 2 hrs), shaking the cage lightly on an orbital shaker (150 rpm, 2 times a week for 2 hrs) and exposure to 37 centigrade temperature (40% humidity, 1 time for 10 mins). For males, social defeat (2 times a week for the time needed to defeat) was used as a social stressor, and social isolation (2 times a week for 12 hrs) was used for females. Social defeat was terminated as soon as intruder (test) mice display signs of social avoidance of the resident (dominant) mouse to avoid any physical wounding. Stressors were pseudorandomized (random order that was kept constant between all cohorts of mice). Before the stress week, all mice underwent a control week during which body weight and core body temperatures were measured at the same times as during stress week.

### Physiological measurements

Animals were weighed daily at the beginning of the dark cycle. Core body temperature was measured using a rectal probe (Kent Scientific, Torrington, CT) inserted 2 cm deep and read as soon as the temperature stabilized. Temperature measurements were taken before the onset of stress and 15-20 minutes after the onset of stress. For control animals, temperature measurements were taken 2 successive times, 15-20 minutes apart, at the same times of day as during stress. Mice were returned to their home cage in between measurements. All temperature differences were averaged per animal before the statistical comparison.

### Adrenal gland isolation

For adrenal gland isolation, mice were anesthetized with a mixture of ketamine and xylazine and transcardially perfused with warm 0.1 M phosphate buffer, followed by warm 4% paraformaldehyde (Electron Microscopy Sciences, Hatfield, PA). Kidneys and adrenals were isolated and postfixed overnight in 4% paraformaldehyde and washed overnight in 0.1 M phosphate buffer. Glands were isolated using a stereomicroscope (Olympus, Tokyo, Japan) and weighed using a precision scale (Mettler Toledo, Columbus, OH). Weight from both glands was averaged and divided by the animal’s weight before the statistical comparison.

### Tail suspension

Mice were suspended by their tail on a bar 55 cm above a platform for 6 minutes and recorded using a high frame rate camera (GoPro, San Mateo, CA), as previously described (Can et al., 2012). Videos were manually scored by experimenters blind to the experimental group being analyzed. Mice were considered immobile if no movement was detected for 3 seconds.

### Open field

Mice were habituated to the room 30 minutes before the behavioral monitoring started. Individual mice were released in a 50×50×50 (all in cm) plexiglass arena with an overhead high frame rate camera (GoPro), and their activity was recorded for 10 minutes. Data was analyzed using MATLAB (Natick, MA)(Zhang et al., 2020).

### Experimental design and data analysis

The experimenters were blind to the genotype or group of animals used during data analysis. Between-subject design was used for physiological measures to allow for the collection of adrenal glands, and within subject design was used for behavioral measures as previous studies suggest no effect of repeated behavioral assays (such as tail suspension) on mouse behavior (Cnops et al. 2022). All data was first compared using 3-way ANOVA (factors: stress, plasticity level and genotype) using DATAtab (Graz, Austria), and followed up using 2-way ANOVA in JASP (JASP Team, 2024) if no 3-way interaction was detected. For 2-way ANOVAs, all data was first tested for interactions between stress and genotype as factors, as well as for the effect of genotype alone if 2-way interaction was not detected. After that, the data was tested for interactions between stress and age, followed by tests for interactions between stress and plasticity, with adolescent WT mice and SynCAM 1 KO mice grouped together into a “high plasticity” group. All data are reported as mean ± SEM or mean ± SD, as indicated, where N represents number of animals used. Target power for all sample sizes was 0.8. In all cases, alpha was set to 0.05.

## Results

### Short bout of unpredictable stress induces robust hyperthermia

As age correlates with neuronal resilience to stress-induced loss of dendritic complexity (Bloss et al., 2010), we hypothesized that the type of plasticity present in the young brain promotes resilience to stress. To test this, we administered a short bout of unpredictable stress to young mice during the 5^th^ postnatal week, when the levels of cortical developmental plasticity are high (Gordon and Stryker, 1996). We did the same to adult (> 2 months old) mice that do not express SynCAM 1 (Synaptic Cell Adhesion Molecule 1), as they display critical period-like cortical plasticity (Ribic et al., 2019). Our stress paradigm was based on the well-established chronic unpredictable mild stress paradigm (Willner et al., 1987), where male and female adult and adolescent wild type (WT), as well as adult SynCAM 1 knock out (KO) mice underwent a pseudo-randomized array of sex-specific and sex-nonspecific stressors over 1 week (Figure 1A). The sex-specific stressors included social isolation for female mice, i.e. group-housed female mice were separated to individual cages for five hours at a time, and social defeat for male mice, i.e. male mice are placed in the resident cage of an aggressor male mouse until they displayed signs of defeat (Golden et al., 2011). The sex nonspecific stressors included restraint, wet bedding, no bedding, cage tilt, cage shaking (at 150 rpm) and heat exposure (37ºC). The length of stress exposure was kept at 1 week to limit it to the peak of critical period plasticity in the cortex (typically the 5^th^ postnatal week)(Gordon and Stryker, 1996; Ribic, 2020)

**Figure 1.**
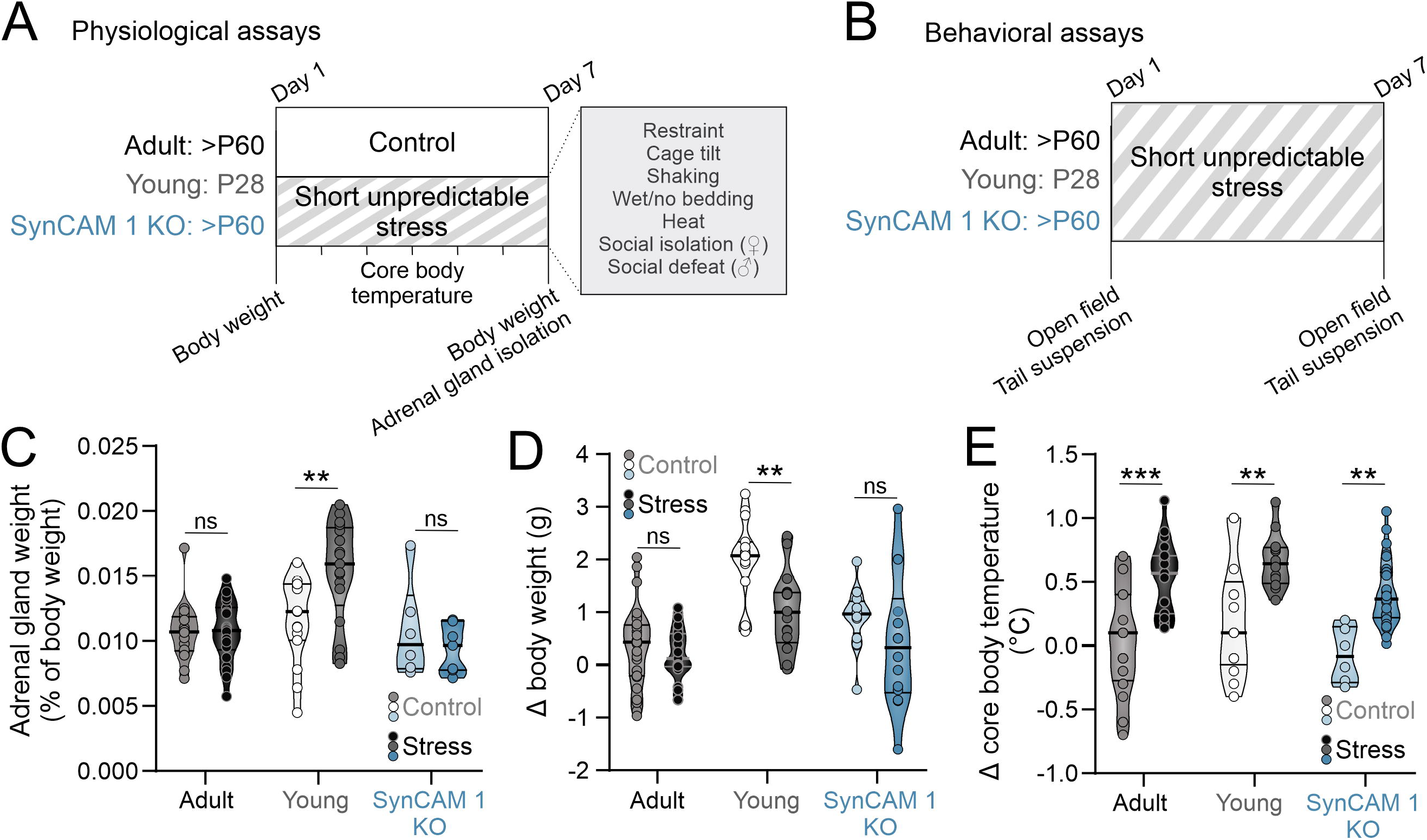
Short unpredictable stress induces robust hyperthermia. **A)** Experimental schematics. Mice of both sexes (adult, postnatal day/P 60 and older; young, P28; adult SynCAM 1 KO) underwent one week of short unpredictable stress with indicated stressors. Control mice of matching ages were subjected to the same measurements but not exposed to the stressors. Body weights and core temperatures were measured daily, and adrenal glands were isolated from both groups of mice after 1 week. **B)** For behavioral assays, mice underwent behavioral assays immediately prior to the first stressor exposure and after the exposure to the last stressor. **C)** Adrenal gland weight was significantly increased in young mice subjected to stress, but not in adult mice. **D)** Short unpredictable abrogated weight gain in young, but not in adult mice. **E)** Short unpredictable stress resulted in robust elevation of core body temperature in all groups of mice. Medians and quartiles of the data are indicated by lines and individual points represent mice.

To test if our stress paradigm indeed induced a physiological stress response, we compared 3 different physiological measures of stress response using a between-subject design (adrenal gland weight, body weight and core body temperature) across different groups of mice: control and stressed WT adults, young WT mice and SynCAM 1 KO mice (Figure 1). For adrenal glands, we normalized their weight to the total body weight of each mouse to account for size differences between young and adult mice, as well as between males and females. A 2-way ANOVA was performed to evaluate the effects of stress and quality of plasticity on adrenal gland weight, where adolescent WT and adult SynCAM 1 KO mice were assigned to a group with “high plasticity” (Tables 1 and 2). We found no significant interaction between stress and genotype (F_(1,80)_=2.095, p=0.152) and no significant effect of genotype alone (p=0.072) on adrenal gland weight. When we tested the interaction between stress and plasticity, we found it to be significant (Table 2). However, further analysis of data revealed that age significantly interacts with stress (p=0.002), and that stress significantly increased adrenal gland weight in adolescent mice of both sexes (Figure 1C; adult mice control vs stress p=0.967, young mice control vs stress p=0.003; Supplementary table 1), but not in adult mice, in agreement with a previous study that demonstrated no changes in adrenal weights after 1 week of unpredictable stress (Luo et al., 2025). As previously reported (Bielohuby et al., 2007), adrenal glands were larger in females than in males across all groups (Supplementary Table 1).

It is well established that different types of stress paradigms in rodents result in reduced body weight (Patterson and Abizaid, 2013). As young mice at baseline are significantly smaller than adult mice of both genotypes (young males=15.95±0.6 g, N=10; young females=13.46±0.23 g, N=13; adult males=22.25±0.42 g, N=11; adult females=17.10±0.44 g, N=12; SynCAM 1 KO males=21.34±0.71 g, N=7; SynCAM1 KO females=17.5±0.85 g, N=6), we compared the difference between body weights at the beginning and at the end of the week in control and stressed mice instead of comparing the body weights themselves (Figure 1D). As with adrenal weights, we found no interaction between stress and genotype (F_(1,116)_, p=0.65) and no effect of genotype alone (p=0.951) on changes in body weight during one week. While the interaction between stress and plasticity was not significant either (Table 2), the interaction between stress and age was (p=0.029), with young stressed mice showing a significant reduction in body weight gain compared to control mice regardless of sex (Figure 1D; adult mice control vs stress p=0.495, young mice control vs stress p=0.004; Supplementary table 1)

Stress induces a robust elevation in core body temperature within 15 minutes, a phenomenon known as stress-induced hyperthermia (SIH)(Vinkers et al., 2008). As our stress paradigm resulted in age-specific adrenal gland weight (Figure 1B) and body weight (Figure 1C) changes, we tested if SIH is present 15-20 minutes after the introduction of each stressor in our paradigm in all groups of mice to ensure that adult mice indeed had a physiological response to our stress paradigm. Rectal readings of core body temperatures can induce stress as well (Zethof et al., 1994), so we compared the differences in temperature (ΔT) before and after the onset of stress to ΔT between two rectal temperature measurements 15-20 minutes apart taken from control mice not exposed to short unpredictable stress (controls; Figure 1D). As expected, we found a robust elevation of core body temperature in stressed mice of both ages and genotypes that was significantly higher than the ΔT of control mice (Tables 1 and 2). While the interaction between stress and genotype was also significant (F_(1,102)_=9.264, p=0.003), post-hoc tests (Tukey) revealed no significant differences between WT and KO control mice, or between WT and KO stressed mice, and no effect of genotype alone (p=0.903). Interactions between stress and age, as well as stress and sex were not significant (p=0.338 and p=0.68, respectively). Altogether, our results demonstrate that a short bout of unpredictable stress induces a robust elevation of core body temperature in all mice, as well as an increase in adrenal gland weight and a reduction in body weight gain in young mice, demonstrating an efficacy of our stress paradigm and indicating age-specific effects of short unpredictable stress on bodily response.

### Stress does not result in depressive-like behaviors in mice with open cortical critical periods

Once we established that our stress paradigm induces a robust physiological response (Figure 1), we tested how mice in whom developmental critical periods of plasticity are still open, namely young wild type (WT) and adult SynCAM 1 KO mice, behaviorally respond to stress compared to adult WT mice. We used two well-established behavioral tests to measure depressive-like and anxiety-like behaviors: tail suspension test (TST) and open field test (OFT; respectively)(Figure 2 and Supplementary Figure 1)(Cryan et al., 2005; Can et al., 2012; Seibenhener and Wooten, 2015).

**Figure 2.**
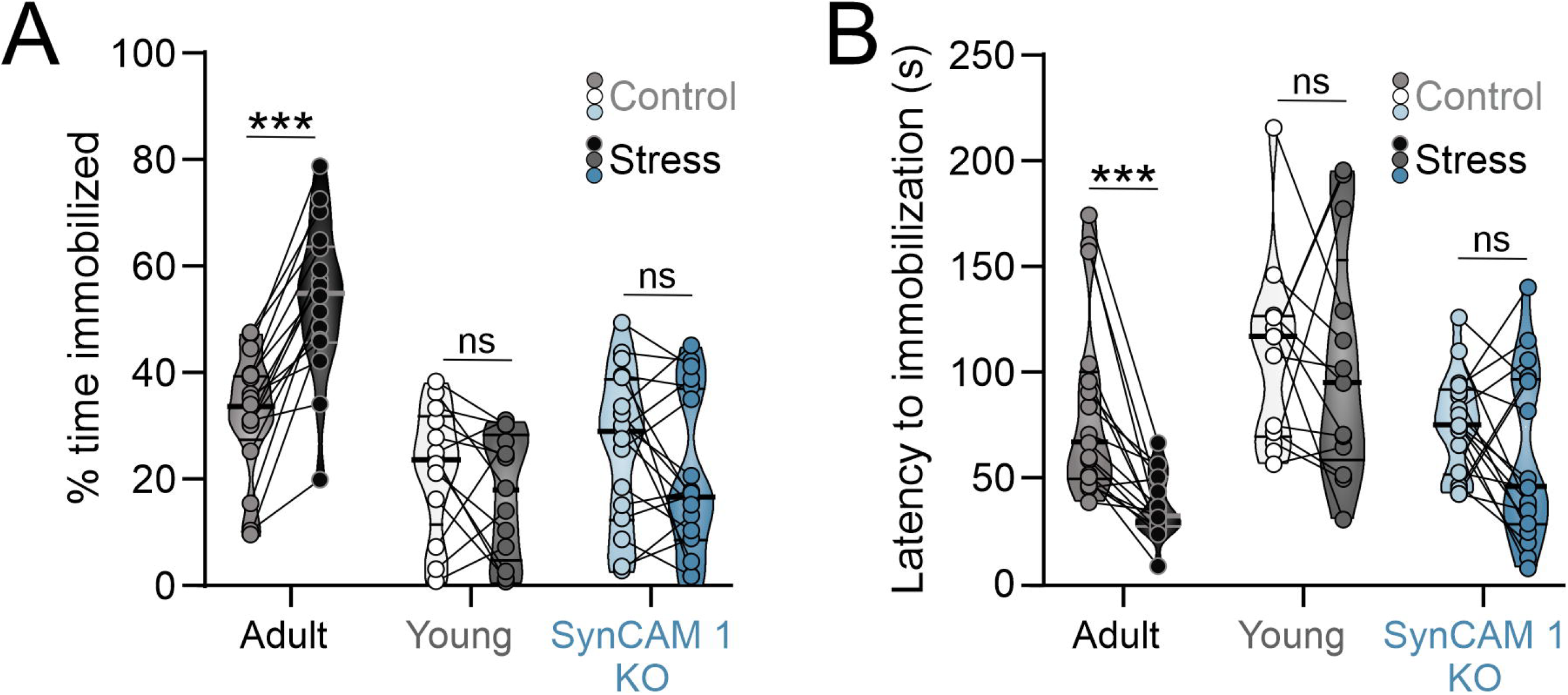
Young mice and adult SynCAM 1 KO mice are resilient to stress-induced increase learned helplessness-like behavior. **A)** Only stressed adult WT mice show increased immobility after stress in tail suspension test and a significant reduction in time lapsed to first immobilization **(B)**. Medians and quartiles of the data are indicated by lines and individual points and lines represent mice.

TST is a measure of learned helplessness-like behavior in rodents that quantifies the fraction of time spent immobilized after exposure to a stressful stimulus, as well as the latency to immobilization. The 2-way interaction between stress and plasticity was significant for the fraction of time spent immobilized (p=0.000005; Table 2) and borderline significant for the latency to first immobilization (p=0.051; Table 2), with post-hoc (Tukey) tests revealing a significant, almost uniform increase in the fraction of time spent immobilized and a significant reduction in time lapsed to first immobilization only in adult WT mice (Figures 2A and 2B; Tables 1 and 2). Interactions between stress and age, as well as stress and sex were not significant (% immobilization: p=0.098 and p=0.55, respectively; latency to first immobilization: p=0.156 and p=0.56, respectively).

Genotype had an effect % immobilization (p=0.017) and post-hoc tests revealed that the difference in immobility between stressed SynCAM 1 KO mice compared to WT mice drives this effect. Genotype had no effect on latency to first immobilization (p=0.18).

To test if our stress paradigm increases anxiety-like behaviors, we compared the preference of mice to stay close to the walls of the open field arena (thigmotaxis, Tables 1 and 2; Supplementary Figure 1A) as well as their locomotor activity (Tables 1 and 2; Supplementary Figure 1B).

Interestingly, we found that short unpredictable stress had no impact on thigmotaxis, while distance travelled in the open field arena was mildly increased in stressed mice from all subgroups, in agreement with previous findings (Sequeira □ Cordero et al., 2019). While the interactions between stress and genotype, as well as stress and age were not significant (p=0.55 and p=0.967, respectively), level of plasticity and genotype alone had significant effects on distance travelled in the open field arena, with SynCAM 1 KO mice traversing the longest distance (Tukey post-hoc; stressed KO vs WT mice, p=0.019), in agreement with increased locomotor activity in these mice (Giza et al., 2013). Locomotor activity in all mouse groups did not correlate with the fraction of time spent immobilized in TST, suggesting that increased locomotor activity of SynCAM 1 KO mice is not the cause of their low immobility in TST assay (Pearson r adult=-0.07, p=0.76; young=0.04, p=0.89; SynCAM 1 KO=-0.02, p=0.91). Altogether, our results demonstrated that short unpredictable stress robustly increases depressive-like behaviors in adult mice, but not in young mice or adult mice with critical period-like cortical plasticity.

## Discussion

Since Hubel and Wiesel reported a temporally restricted window of plasticity in the kitten cortex during which neurons robustly respond to changes in sensory input (Hubel and Wiesel, 1970), many studies have attempted to restore that type of plasticity to the adult brain aiming to mitigate the damage caused by a myriad of stressors (Ribic, 2020; Hensch and Quinlan, 2018; Levelt and Hübener, 2012). In support of this longstanding notion, antidepressants can restore juvenile-like plasticity to the adult cortex (Maya Vetencourt et al., 2008), but it is still unclear if they promote recovery from stress through changing the quality of plasticity in the adult brain. Our study attempted to shed more light on this issue by testing if short unpredictable stress results in depressive-like behavior in adolescent mice in whom the cortical window of plasticity is still open, as well as in adult mice in whom the plasticity windows are extended into adulthood through deletion of SynCAM 1 (Ribic et al., 2019). We found that this is indeed the case, supporting that interventions that change the quality of plasticity in the adult brain can facilitate stress resilience (Nestler and Russo, 2024; McEwen, 2016; Fuchs et al., 2004).

The duration and type of stressors, along with age and sex of animals, can have a significant impact on physiological measures of stress, as evident in our results as well (Luo et al., 2025; Monteiro et al., 2015). Our stress paradigm abrogated weight gain in young mice, but not in adults, indicating the sensitivity of bodily homeostasis to stress in maturing mice. Similarly, adrenal gland weight was increased in young mice only, suggesting that the hypothalamic-pituitary-adrenal (HPA) axis is uniquely sensitive to stress in young mice and may mediate the abrogated weight gain during stress in these mice. While our stress paradigm did not impact weight gain and adrenal weights of adult mice, all 3 groups of mice displayed robust stress-induced hyperthermia, indicating that our protocol was stressful to them. In agreement, adult mice displayed an increase in depressive-like behaviors as measured with tail suspension test (TST). Interestingly, adolescent mice and adult SynCAM 1 KO mice did not show any changes in immobilization after stress, in support of the notion that juvenile/adolescent-like plasticity promotes resilience to stress. The increase in immobilization in WT mice was almost uniform, while young WT and adult SynCAM 1 KO mice displayed significant variability that was not sex-dependent, suggesting a gradation of resilience in these groups of mice. Future studies need to address if the variability in depressive-like measures after stress in young WT and adult SynCAM 1 KO mice correlates with the variability in the level of cortical plasticity. While it is possible that increased locomotor activity of SynCAM 1 KO mice results in reduced immobilization during TST, we found no correlation between immobility and locomotor activity in the open field arena, indicating that young mice and adult SynCAM 1 mice indeed are resilient to short unpredictable stress. Based on our and other studies (Hu et al., 2025), Future studies can now address the impact of longer stress duration or more severe stressors on the behavior of adolescent and adult SynCAM 1 KO mice, and whether juvenile/adult-like plasticity provides a lasting protection from stress.

Unlike immobilization, locomotor activity was mildly impacted by stress, indicated by increased distance travelled in the open field arena in all three groups of mice. While post-hoc comparisons of controlled and stressed mice within different subgroups were not statistically significant, our results are in line with previous research that demonstrated stress-associated hyperlocomotion (Sequeira□Cordero et al., 2019). Distance travelled positively correlates with corticosterone levels in mice head-fixed above a treadmill, which itself is stressful (Juczewski et al., 2020), suggesting that increased locomotion is a stress response. Locomotor activity is traditionally classified as anxiety-like behavior, indicating that juvenile/adolescent-like plasticity may not promote resilience to anxiety after stress. However, future studies need to address this issue using paradigms that can assess anxiety-like phenotype independent of locomotion, like novelty suppressed feeding (NSF)(Koskinen and Hovatta, 2023).

While the type of plasticity in our study correlates with resilience to short unpredictable stress, it is possible that the resilience of SynCAM 1 KO mice is due to another mechanism.

SynCAM 1 gene expression is not restricted to the brain (Fujita et al., 2006) and, while SynCAM 1 protein is localized to the membrane, it can trigger multiple signaling cascades that promote neuronal resilience to stress (Cheadle and Biederer, 2012; Murakami et al., 2014). However, we find this unlikely, as SynCAM 1 KO mice display overall accelerated learning in spatial tasks and juvenile-like responses to noxious stimuli (Park et al., 2016; Robbins et al., 2010). SynCAM 1 is a potent plasticity brake in the adult brain and even a transient knock-down of its expression can restore juvenile/adolescent like plasticity in the adult primary visual cortex (Ribic et al., 2019). Our study is hence in line with the notion that critical period-like plasticity is protective of stress and provides a rationale for further inquiries into the link between resilience and different types of plasticity.

Our study is limited in its scope, but our findings unambiguously demonstrate the efficacy of short unpredictable stress in inducing depressive-like behaviors in adult mice. The short duration and ease of administering our protocol will undoubtedly facilitate further studies of short unpredictable stress and its long-term effects on brain and behavior. Importantly, our study provides indirect evidence that heightened brain plasticity promotes resilience to stress, and provides a rationale for future, more in-depth mechanistic inquiries into the relationship between plasticity and stress.

## Author Contributions

AR and RW conceived and designed the study. RW, CVDO, ENL, SCF, GLO and AR collected and analyzed the data. AR wrote the manuscript with input from all authors.

## Acknowledgments

The authors would like to thank the Ribic Lab for critical reading of the manuscript.

## Data Availability Statement

The datasets generated and analyzed in this study will be provided upon reasonable request.

## Figure Legends

**Supplementary Figure 1. Short unpredictable stress has no impact on anxiety-like behaviors. A)** Short unpredictable stress did not impact thigmotaxis in open field test. **B)** While stressed mice from all groups travelled more in the open field arena, within-group post-hoc comparisons were not statistically significant. Medians and quartiles of the data are indicated by lines and individual points and lines represent mice.

