## Supplemental table 1 for "Critical Period Plasticity is Associated with Resilience to Short Unpredictable Stress"

**Supplementary Table 1.** *Descriptive statistics of adrenal gland weights and Δ body weights by sex*

| Measure | Animal group | Sex | Control | | | Stress | | |
| --- | --- | --- | --- | --- | --- | --- | --- | --- |
|  |  |  | *M* | *SD* | *N* | *M* | *SD* | N |
| Adrenal gland weight  (% of body weight) | Adult WT (low plasticity) | M | 0.008 | 0.009 | 3 | 0.01 | 0.002 | 12 |
|  |  | F | 0.012 | 0.002 | 11 | 0.012 | 0.002 | 10 |
|  | Young WT (high plasticity) | M | 0.006 | 0.0017 | 3 | 0.011 | 0.003 | 6 |
|  |  | F | 0.012 | 0.002 | 7 | 0.018 | 0.002 | 11 |
|  | Adult SynCAM 1 KO (high plasticity) | M | 0.009 | 0.0016 | 3 | 0.009 | 0.002 | 4 |
|  |  | F | 0.013 | 0.004 | 3 | 0.011 | 0.001 | 3 |
| Δ body weight | Adult WT (low plasticity) | M | 0.081 | 0.550 | 14 | 0.284 | 0.403 | 11 |
|  |  | F | 0.234 | 0.693 | 20 | 0.13 | 0.590 | 11 |
|  | Young WT (high plasticity) | M | 2.190 | 0.87 | 7 | 0.813 | 0.614 | 6 |
|  |  | F | 1.738 | 0.664 | 8 | 1.042 | 0.795 | 10 |
|  | Adult SynCAM 1 KO (high plasticity) | M | 1.005 | 0.608 | 10 | 0.646 | 1.289 | 8 |
|  |  | F | 0.752 | 0.402 | 6 | 0.137 | 1.247 | 6 |
