## Supplementary material for "Critical Period Plasticity is Associated with Resilience to Short Unpredictable Stress": Table 1

**Table 1.** *Descriptive statistics for physiological and behavioral measures obtained*

| Measure | Animal group | Control | | | Stress | | |
| --- | --- | --- | --- | --- | --- | --- | --- |
|  |  | *M* | *SD* | *N* | *M* | *SD* | N |
| Adrenal gland weight (% of body weight) | Adult WT (low plasticity) | 0.011 | 0.002 | 17 | 0.011 | 0.002 | 22 |
|  | Young WT (high plasticity) | 0.012 | 0.003 | 15 | 0.015 | 0.004 | 17 |
|  | Adult SynCAM 1 KO (high plasticity) | 0.011 | 0.004 | 6 | 0.009 | 0.002 | 7 |
| Δ body weight | Adult WT (low plasticity) | 0.329 | 0.733 | 33 | 0.208 | 0.486 | 25 |
|  | Young WT (high plasticity) | 1.949 | 0.774 | 15 | 0.996 | 0.712 | 17 |
|  | Adult SynCAM 1 KO (high plasticity) | 0.910 | 0.540 | 16 | 0.428 | 1.249 | 14 |
| Δ temperatures | Adult WT (low plasticity) | 0.038 | 0.412 | 24 | 0.538 | 0.282 | 19 |
|  | Young WT (high plasticity) | 0.215 | 0.456 | 13 | 0.665 | 0.218 | 12 |
|  | Adult SynCAM 1 KO (high plasticity) | -0.072 | 0.217 | 6 | 0.406 | 0.251 | 32 |
| % immobilization^#^ | Adult WT (low plasticity) | 31.786 | 11.508 | 17 | 53.729 | 14.419 | 17 |
|  | Young WT (high plasticity) | 21.741 | 12.155 | 13 | 16.752 | 11.595 | 13 |
|  | Adult SynCAM 1 KO (high plasticity) | 26.657 | 13.965 | 18 | 21.003 | 14.614 | 18 |
| Latency to immobilization^#^ | Adult WT (low plasticity) | 82.706 | 43.066 | 17 | 37.176 | 14.972 | 17 |
|  | Young WT (high plasticity) | 108.538 | 43.395 | 13 | 103.462 | 55.583 | 13 |
|  | Adult SynCAM 1 KO  (high plasticity) | 75.722 | 23.539 | 18 | 60.056 | 39.740 | 18 |
| Thigmotaxis^#^ | Adult WT (low plasticity) | 0.828 | 0.070 | 29 | 0.818 | 0.074 | 29 |
|  | Young WT (high plasticity) | 0.759 | 0.035 | 9 | 0.783 | 0.059 | 9 |
|  | Adult SynCAM 1 KO (high plasticity) | 0.812 | 0.088 | 27 | 0.808 | 0.082 | 27 |
| Distance travelled^#^ | Adult WT (low plasticity) | 24.258 | 5.264 | 29 | 26.389 | 8.243 | 29 |
|  | Young WT (high plasticity) | 29.778 | 5.864 | 9 | 32.663 | 5.628 | 9 |
|  | Adult SynCAM 1 KO (high plasticity) | 30.053 | 9.957 | 27 | 34.138 | 10.121 | 27 |

^#^Within-subject design
