## Supplementary material for "Critical Period Plasticity is Associated with Resilience to Short Unpredictable Stress": Table 2

**Table 2.** *Statistical comparison of data in Figures 1 and 2*

|  | **Factors** | **Sum of Squares** | **df** | **Mean Square** | **F** | **p** | **η²** |
| --- | --- | --- | --- | --- | --- | --- | --- |
| Adrenal gland weight | Stress | 2.489×10^-5^ | 1 | 2.489×10^-5^ | 2.338 | 0.130 | 0.026 |
|  | Plasticity | 6.405×10^-5^ | 1 | 6.405×10^-5^ | 6.017 | 0.016 | 0.066 |
|  | Stress x Plasticity | 2.808×10^-5^ | 1 | 2.808×10^-5^ | 2.638 | 0.108 | 0.029 |
|  | Residuals | 8.515×10^-4^ | 80 | 1.064×10^-5^ |  |  |  |
| Δ Body Weights | Stress | 4.677 | 1 | 4.677 | 7.247 | 0.008 | 0.046 |
|  | Plasticity | 19.367 | 1 | 19.367 | 30.008 | 2.534×10^-7^ | 0.191 |
|  | Stress x Plasticity | 2.258 | 1 | 2.258 | 3.498 | 0.064 | 0.022 |
|  | Residuals | 74.865 | 116 | 0.645 |  |  |  |
| Δ Temperatures | Stress | 3.283 | 1 | 3.283 | 27.135 | 9.913×10^-7^ | 0.201 |
|  | Plasticity | 0.227 | 1 | 0.227 | 1.876 | 0.174 | 0.014 |
|  | Stress x Plasticity | 0.491 | 1 | 0.491 | 4.060 | 0.047 | 0.030 |
|  | Residuals | 12.339 | 102 | 0.121 |  |  |  |
| % immobilization | Stress | 1506.843 | 1 | 1506.843 | 8.613 | 0.004 | 0.048 |
|  | Plasticity | 9546.112 | 1 | 9546.112 | 54.562 | 6.656×10^-11^ | 0.306 |
|  | Stress x Plasticity | 4096.759 | 1 | 4096.759 | 23.416 | 5.233×10^-6^ | 0.131 |
|  | Residuals | 16096.159 | 92 | 174.958 |  |  |  |
| Latency to immobilization | Stress | 17682.798 | 1 | 17682.798 | 10.735 | 0.001 | 0.094 |
|  | Plasticity | 12574.108 | 1 | 12574.108 | 7.633 | 0.007 | 0.067 |
|  | Stress x Plasticity | 6459.798 | 1 | 6459.798 | 3.921 | 0.051 | 0.034 |
|  | Residuals | 151549.677 | 92 | 1647.279 |  |  |  |
| Thigmotaxis | Stress | 4.504×10^-4^ | 1 | 4.504×10^-4^ | 0.078 | 0.780 | 6.047×10^-4^ |
|  | Plasticity | 0.017 | 1 | 0.017 | 2.904 | 0.091 | 0.022 |
|  | Stress x Plasticity | 0.001 | 1 | 0.001 | 0.235 | 0.629 | 0.002 |
|  | Residuals | 0.726 | 126 | 0.006 |  |  |  |
| Distance travelled in open field arena | Stress | 281.067 | 1 | 281.067 | 4.180 | 0.043 | 0.028 |
|  | Plasticity | 1379.307 | 1 | 1379.307 | 20.513 | 1.356×10^-5^ | 0.136 |
|  | Stress x Plasticity | 21.963 | 1 | 21.963 | 0.327 | 0.569 | 0.002 |
|  | Residuals | 8472.303 | 126 | 67.240 |  |  |  |

*Note.*  Type III Sum of Squares
