## Supplementary figures and images for "Critical Period Plasticity is Associated with Resilience to Short Unpredictable Stress"

### Supplemental Figure 1

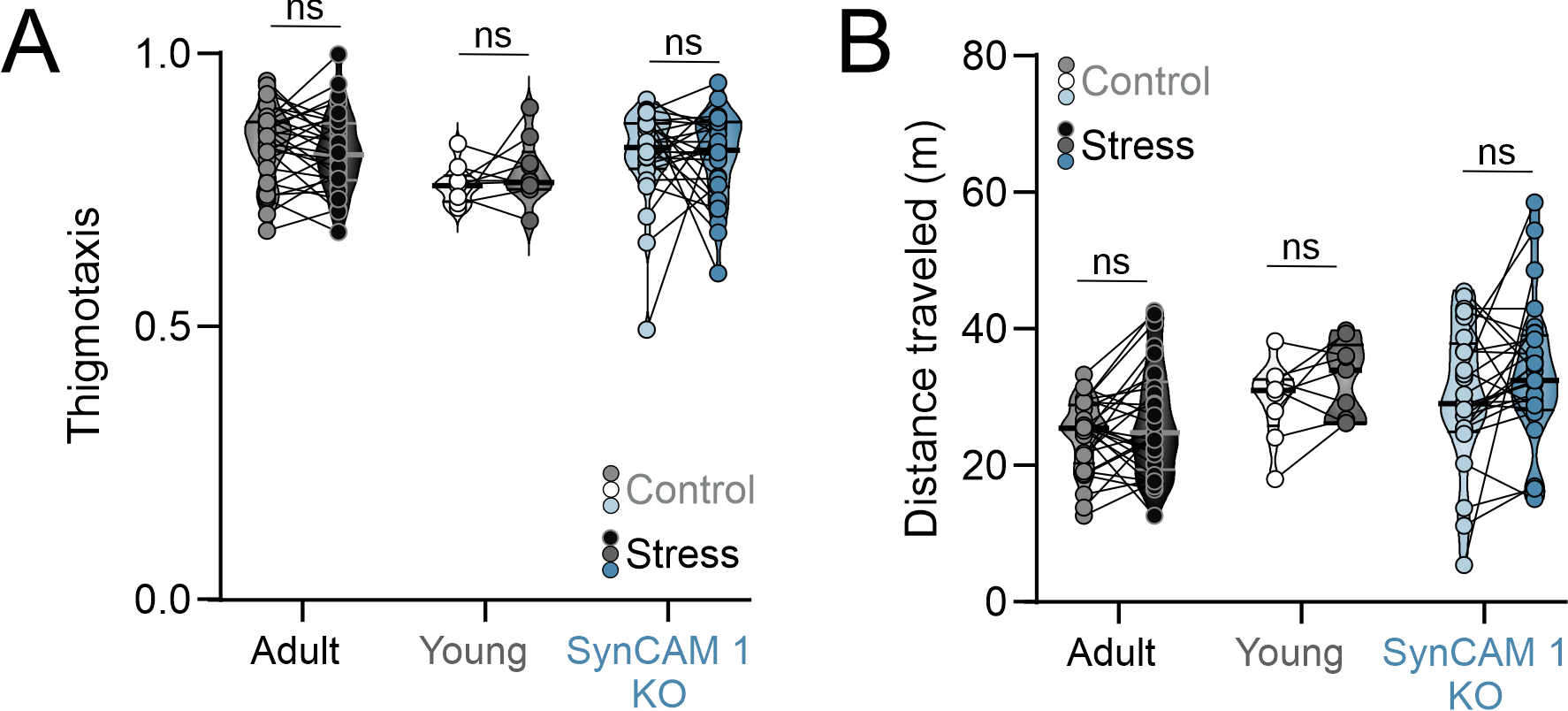
